# Molecular Basis for the Evolved Instability of a Human G-Protein Coupled Receptor

**DOI:** 10.1101/2019.12.20.884718

**Authors:** Laura M. Chamness, Nathan B. Zelt, Charles P. Kuntz, Brian J. Bender, Wesley D. Penn, Joshua J. Ziarek, Jens Meiler, Jonathan P. Schlebach

**Affiliations:** Department of Chemistry, Indiana University, Bloomington, Indiana, USA; Department of Molecular and Cellular Biochemistry, Indiana University, Bloomington, Indiana, USA; Department of Chemistry, Vanderbilt University, Nashville, TN, USA; Institut for Drug Development, Leipzig University, Leipzig, SAC, Germany

**Keywords:** GnRHR, GPCR, Membrane Protein Folding, Proteostasis, Epistasis

## Abstract

Membrane proteins are prone to misfolding and degradation. This is particularly true for mammalian forms of the gonadotropin-releasing hormone receptor (GnRHR). Though they function at the plasma membrane, mammalian GnRHRs tend to accumulate within the secretory pathway. Their apparent instability is believed to have evolved in response to selection for attenuated GnRHR activity. Nevertheless, the structural basis of this adaptation remains unclear. We find that this adaptation coincides with a C-terminal truncation and an increase in the polarity of its transmembrane (TM) domains. This enhanced polarity compromises the translocon-mediated cotranslational folding of two TM domains. Moreover, replacing a conserved polar residue in TM6 with an ancestral hydrophobic residue partially restores GnRHR expression with minimal impact on function. An evolutionary analysis suggests variations in the polarity of this residue are associated with reproductive differences. Our findings suggest the marginal energetics of cotranslational folding can be exploited to tune membrane protein fitness.

## Introduction

Proteins must continually sample new mutations that modify their conformational equilibria in order to maximize their evolutionary fitness (*1*). Most mutations destabilize native protein structures, and evolutionary pathways are limited to those in which the protein retains sufficient conformational stability and activity with each successive mutation (*2*). Like water-soluble proteins, the sequences of integral membrane proteins (MPs) are also constrained by folding energetics (*3*). However, their transmembrane (TM) domains must also retain sufficient hydrophobicity to partition into the membrane, which further constrains their evolutionary sequence space (*3*). In recent investigations of the sequence constraints within the class A G-protein coupled receptor (GPCR) rhodopsin, we found its expression to be highly sensitive to mutations within a marginally hydrophobic TM domain (*4*). Moreover, the natural sequence of this TM domain appears to be more polar than is necessary to support function, and its expression can be enhanced by functionally-neutral hydrophobic substitutions (*4*). As a result of this instability, much of the nascent protein is retained in the endoplasmic reticulum (ER) (*4*). As is true for water-soluble proteins, these observations suggest that naturally evolved MPs tend to be metastable, and have not evolved to maximize the efficiency of protein biogenesis. Nevertheless, it is unclear whether this metastability provides an evolutionary benefit, or if it is simply an emergent property that stems from the evolutionary process itself.

A series of previous investigations of the gonadotropin releasing hormone receptor (GnRHR), another class A GPCR, revealed that the mammalian forms of these receptors exhibit a heightened tendency to misfold and accumulate within the ER (*5*). GnRHR plays a critical role in steroidogenesis, and its functional expression at the plasma membrane is critical for reproductive fitness (*6*, *7*). Variations within the *GnRHR* gene have been associated with shifts in litter size and length of the luteal phase (*8*–*11*). Additionally, numerous loss of function mutations in human *GnRHR* have been found to cause hypogonadotropic hypogonadism (HH), which is characterized by infertility and loss of gonadal function (*6*). GnRHRs found in fish, which produce many offspring, appear to exhibit robust plasma membrane expression (PME) relative to those found in mammals, which have far fewer offspring (*7*, *9*). Based on various observations, it has been suggested that the reproductive selection pressures increased the relative fitness of mammals expressing less stable GnRHR variants with diminished PME (*10*, *12*). It has also been speculated that the pool of immature GnRHRs in the ER may provide a regulatory benefit, as modifications to the proteostasis network can alter the flux of mature protein through the secretory pathway (*7*, *9*, *13*). Thus, it appears as though nature may have exploited the instability of GnRHRs in order to tune their evolutionary fitness. Nevertheless, the nature of the conformational modifications involved in these evolutionary adaptations remain poorly understood.

To evaluate the evolutionary sequence modifications that coincide with the proteostatic divergence between the mammalian and non-mammalian GnRHRs, we first compiled and analyzed the sequences of known type I GnRHRs. Consistent with previous observations, we first show that a C-terminal truncation within the mammalian receptors appears to significantly contribute to the attenuation of the mammalian receptor PME. However, multiple sequence alignments also reveal that the transmembrane (TM) helices of mammalian GnRHRs are considerably more polar than those of non-mammalian GnRHRs. We show that this enhanced polarity compromises the translocon-mediated membrane integration of two of its seven TM domains, which demonstrates that some of these sequence modifications compromise the fidelity of cotranslational folding. Structural models of these receptors suggest two of the polar substitutions that compromise translocon-mediated membrane integration of the nascent chain occur at surface residues that are projected into the membrane core. Moreover, we show that re-introducing the ancestral hydrophobic side chain at one of these positions partially restores the PME of human GnRHR with minimal impact on receptor activation. Finally, we show that natural variations in the polarity of this residues among mammalian GnRHRs are associated with dramatic variations in litter size. Together, these findings provide new evidence that evolution has exploited the marginal cotranslational folding energetics of GnRHR in order to tune its fitness. These observations suggest the instability of natural proteins provides an additional avenue for evolutionary adaptation.

## Results

### Cellular Expression of Natural GnRHRs

Various lines of evidence suggest the selection for reduced GnRH signaling in higher mammals produced GnRHRs with diminished conformational stability and attenuated plasma membrane expression (PME) (*5*, *14*). Nevertheless, most previous studies relied on activity as a proxy for PME (*7*, *15*, *16*). To more directly probe differences in GnRHR expression, we employed immunostaining in conjunction with flow cytometry to quantitatively characterize the expression of three previously characterized GnRHRs (human, mouse, and catfish). Briefly, each of these receptors was transiently expressed in HEK293T cells prior to labeling plasma membrane and intracellular GnRHRs with distinct fluorescent antibodies, as previously described (*17*). Cellular fluorescence profiles were then analyzed by flow cytometry. A comparison of the distribution of single-cell fluorescence profiles reveals that larger proportions of the expressed mouse GnRHR (*Mus musculus*, mGnRHR) and catfish GnRHRs (*Clarius gariepinus*, cGnRHR) accumulate at the plasma membrane relative to human GnRHR (*Homo sapiens*, hGnRHR, Figure 1). The mean fluorescence intensity associated with the surface immunostaining of hGnRHR at the plasma membrane is 21.5 ± 6.0 fold lower than that of mGnRHR and 92.0 ± 17.9 fold lower than that of cGnRHR. Overall, the total cellular expression of hGnRHR was 2.71 ± 0.04 fold lower than that of mGnRHR and 2.04 ± 0.35 fold lower than cGnRHR. These results show for the first time that cGnRHR exhibits robust expression and trafficking relative to the mammalian receptors under equivalent conditions. Nevertheless, the nature of the structural modifications responsible for this apparent proteostatic divergence remains unclear.

**Figure 1.**
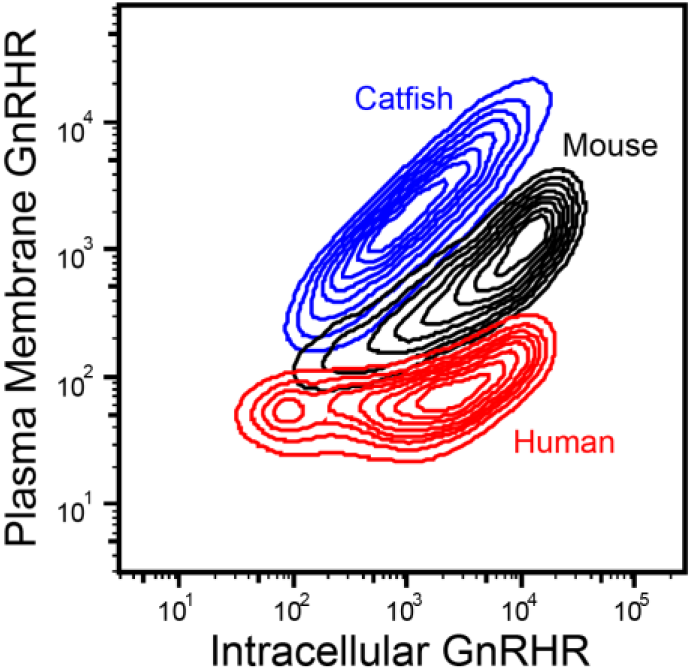
Cellular trafficking of GnRHR variants in HEK293T cells. Human (red), mouse (black), and catfish (blue) GnRHRs were transiently expressed in HEK293T cells, and the relative abundance of plasma membrane GnRHR and intracellular GnRHR was analyzed by flow cytometry. Contour plots show the distribution of cellular fluorescence intensities associated with immunostaining of plasma membrane (y-coordinate) and intracellular (x-coordinate) GnRHRs.

### Impact of the C-terminal Tail in GnRHR Expression

Evolutionary adaptations in mammalian GnRHRs coincided with a variety of sequence modifications. Most strikingly, mammalian GnRHRs feature a C-terminal deletion of a disordered loop as well as a conserved amphipathic helix (helix 8, H8) that contains two palmitoylation sites (*14*, *18*, *19*). Fusing the C-terminal domain of cGnRHR to the C-terminus hGnRHR was previously shown to enhance the activity of the human receptor (*5*). Nevertheless, the extent to which the structural elements within the C-terminal region impact the PME remains unclear. We therefore assessed the effects of various C-terminal modifications on the PME of cGnRHR. To determine whether C-terminal palmitoylation impacts PME, we first characterized a double mutant of cGnRHR that lacks its two C-terminal palmitoylation sites (C339A, C341A). Removal of these palmitoylation sites has minimal effect on the PME of cGnRHR (Figure S1), which suggests the loss of these modifications is not responsible for the attenuated PME of the mammalian receptors. To determine whether the disordered portion of the C-terminal tail impacts PME, we next characterized a cGnRHR variant with a deletion downstream of H8 (Δ352-379). Truncation of these residues reduces the PME of cGnRHR 2.0 ± 0.2 fold, (Figure S1), which suggests these residues are important for efficient expression. Finally, to determine whether H8 impacts PME, we characterized a cGnRHR variant lacking the entire C-terminal tail (Δ329-379), which mimics the deletion found in mammalian receptors (*18*). This truncation reduces the PME of cGnRHR 19.4 ± 0.9 fold (Figure S1), which demonstrates that the truncation of the C-terminal domain is primarily responsible for the diminished PME of mammalian GnRHRs. Nevertheless, the PME of this truncated cGnRHR variant is still 4.2 ± 1.1 fold higher than that of hGnRHR. Therefore, it is likely that additional sequence modifications have helped to tune the PME of mammalian GnRHRs.

### Impact of Sequence Variations on the Topological Energetics of GnRHR

Previous investigations have concluded that evolutionary modifications of PME arise from variations in the conformational stability of GnRHR (*5*, *10*). Variations in conformational stability may impact the propensity of the receptor to misfold during translocon-mediated cotranslational folding (stage I folding) and/ or during post-translational folding (stage II folding) (*20*). The efficiency of stage I folding primarily depends on the hydrophobicity of TM domains, which dictates the energetics of their translocon-mediated membrane integration (*21*). To assess whether evolutionary adaptations may impact the fidelity of stage I folding, we analyzed the sequences of cGnRHR and hGnRHR using a knowledge-based algorithm that predicts the free energy difference associated with the transfer of nascent TM domains from the translocon to the ER membrane (ΔG predictor) (*22*). A scan of the cGnRHR sequence reveals that six of its seven TM domains have pronounced energetic minima, four of which have negative transfer free energies (Figure S2). This observation suggests that most TM domains within cGnRHR are sufficiently hydrophobic to undergo efficient translocon-mediated membrane integration. By comparison, only two of the seven TM domains within hGnRHR have negative transfer free energies (Figure S2), which suggests this protein may be more prone to the formation of topological defects during stage I folding.

To determine whether these differences are reflective of a wider evolutionary trend, we used the ΔG predictor to scan the sequences of a total 59 known GnRHRs (Table S1). Projection of the average predicted transfer free energies across the seven TM domains of each receptor onto a phylogenetic tree reveals stark contrasts in the topological energetics of mammalian and non-mammalian GnRHRs (Figure 2). The average predicted transfer free energies are significantly higher for mammalian GnRHRs relative to those of the non-mammalian receptors (Figure 3A, Mann-Whitney *p* = 5 × 10^−14^). A comparison of the distribution of predicted transfer free energies for individual domains reveals that evolutionary adaptations resulted in particularly stark increases in the polarity of TM2 and TM6 (Figure 3B). It is unclear how sequence modifications within these domains may have impacted the energetics of post-translational folding reactions or functional signaling. Nevertheless, heightened predicted transfer free energies of mammalian TM2 and TM6 suggest the modifications within these regions could potentially compromise the efficiency of the cotranslational folding of mammalian GnRHRs.

**Figure 2.**
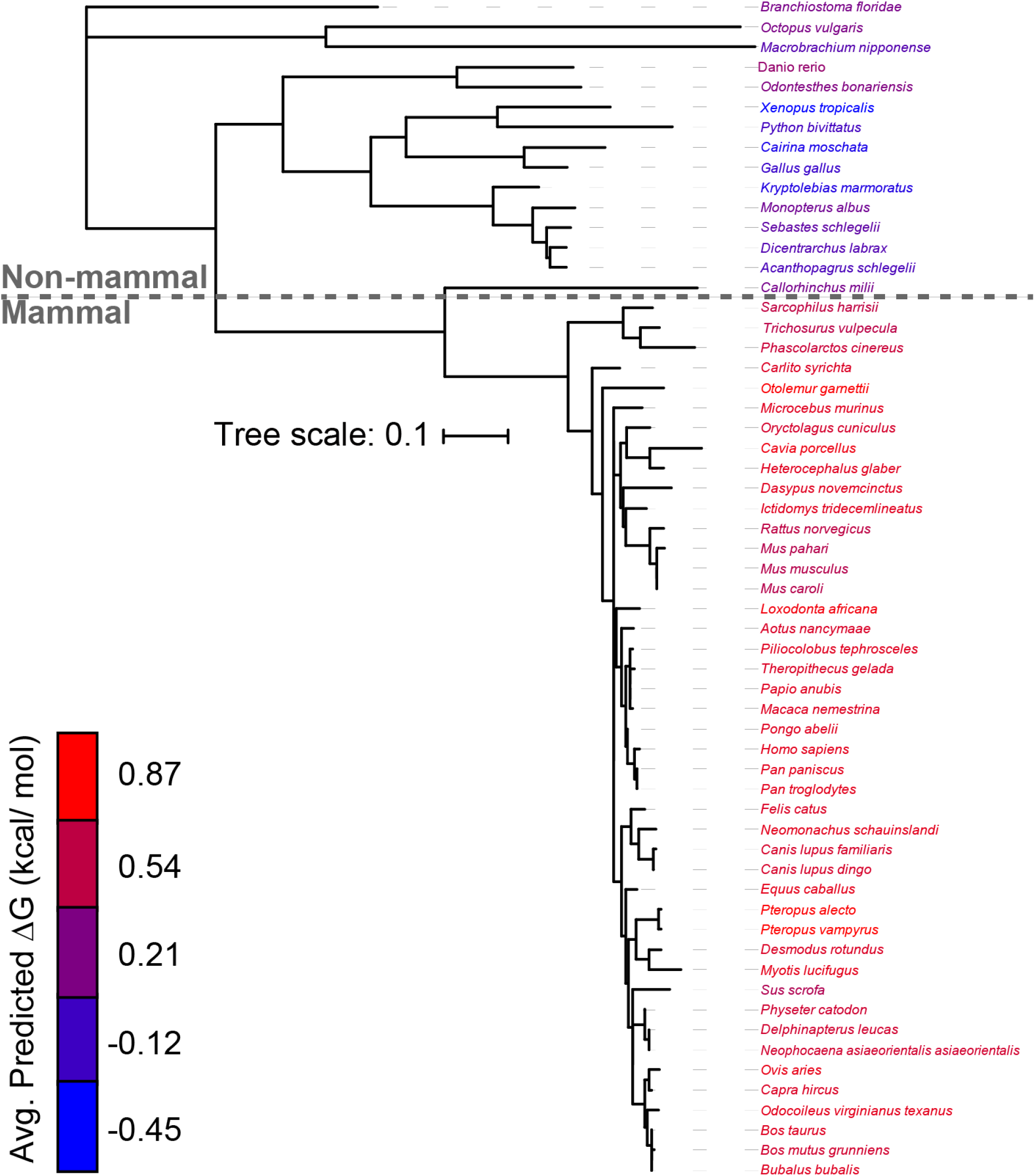
Evolutionary divergence of the topological energetics of GnRHR. The predicted transfer free energy associated with the translocon-mediated membrane integration of each TM domain within 59 known GnRHRs was calculated using the ΔG predictor (*22*), and the average value for the seven TM domains within each receptor was projected onto a phylogenetic tree. The names of each species within the phylogenetic tree are colored according to the average ΔG value for the seven TM domains within the corresponding receptor. The phylogenetic tree was constructed using ML after MUSCLE alignment using MEGA7 software, and the branch lengths represent number of substitutions per site. A length unit of 0.1 substitutions per site corresponds to 10% divergence. The *Clarius gariepinus* (catfish) receptor is annotated as a type II GnRHR, and was therefore excluded from this analysis (see Methods).

**Figure 3.**
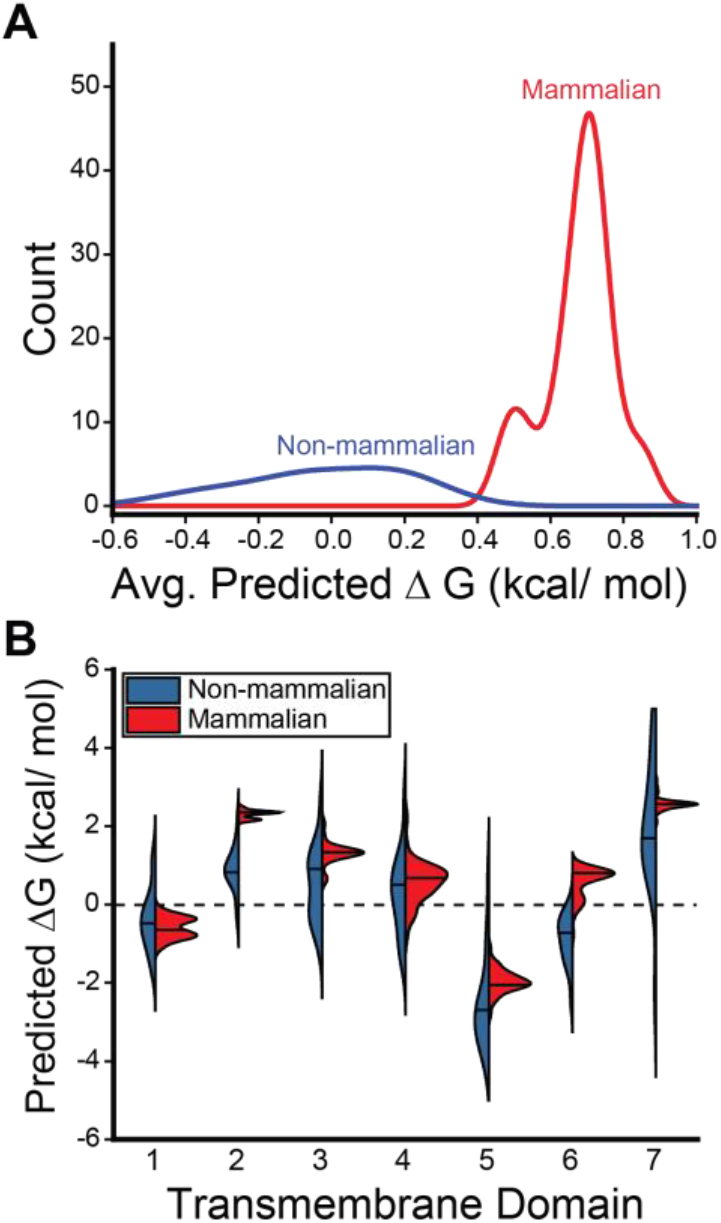
Comparison of the topological energetics of mammalian and non-mammalian GnRHRs. The distribution of predicted transfer free energies associated with the translocon-mediated membrane integration of the TM domains within 59 known GnRHRs are shown. A) A histogram depicts the distribution of the average transfer free energies across the seven TM domains of the non-mammalian (blue) and mammalian (red) GnRHRs. B) A series of violin plots depict the distribution of predicted transfer free energies for each individual TM domain found within the non-mammalian (blue) and mammalian (red) receptors. The position of the median value is indicated by a horizontal line within each distribution. The shapes of the histograms and violins were generated using a kernel smoothing function.

### Impact of Polar Substitutions on the Cotranslational Folding of TMs 2 & 6

To further explore these TM domains, we constructed logo plots that depict the most common amino acids found at each position within TM2 and TM6 of mammalian and non-mammalian GnRHRs. The sequences of the non-mammalian TM domains are more diverse than those of the mammalian receptor (Figure 4 A & B), which reflects the increased evolutionary distance between the non-mammalian sequences (Figure 2). Nevertheless, a comparison of the most common amino acids at each position reveals the heightened transfer free energies of mammalian GnRHRs primarily arise from three polar substitutions in TM2 and two polar substitutions in TM6 (Figure 4 A & B). To assess the impact of these substitutions on the fidelity of stage I folding, we compared the translocon-mediated membrane integration of the consensus versions of the mammalian and non-mammalian TM domains. Briefly, a series of chimeric leader peptidase (Lep) proteins containing each TM domain of interest was produced by *in vitro* translation in canine rough microsomes. The membrane integration efficiency of each TM domain can then be inferred from the glycosylation state of Lep; membrane integration of the TM domain results in a single glycosylation whereas passage into the lumen generates two glycosyl modifications (Figure 4C). The Lep protein containing the non-mammalian TM2 is produced as a mix of both glycoforms (Figure 4D). In contrast, the doubly glycosylated form predominates for the Lep protein containing the mammalian TM2 (Figure 4D), which demonstrates that the increased polarity of the mammalian TM2 compromises its recognition by the translocon. Similarly, the membrane integration of the non-mammalian form of TM6 appears to be significantly more efficient than that of the corresponding mammalian form (Figure 4D). The apparent transfer free energies associated with the translocon-mediated membrane integration of these helices slightly deviate from the predicted values (Table S2). Nevertheless, these predictions correctly predicted the manner in which these mutations should impact the efficiency of membrane integration. Thus, consistent with computational predictions (Figure 3B), these results show that the increased polarity of TM2 and TM6 of mammalian GnRHRs decreases the efficiency of their translocon-mediated membrane integration.

**Figure 4.**
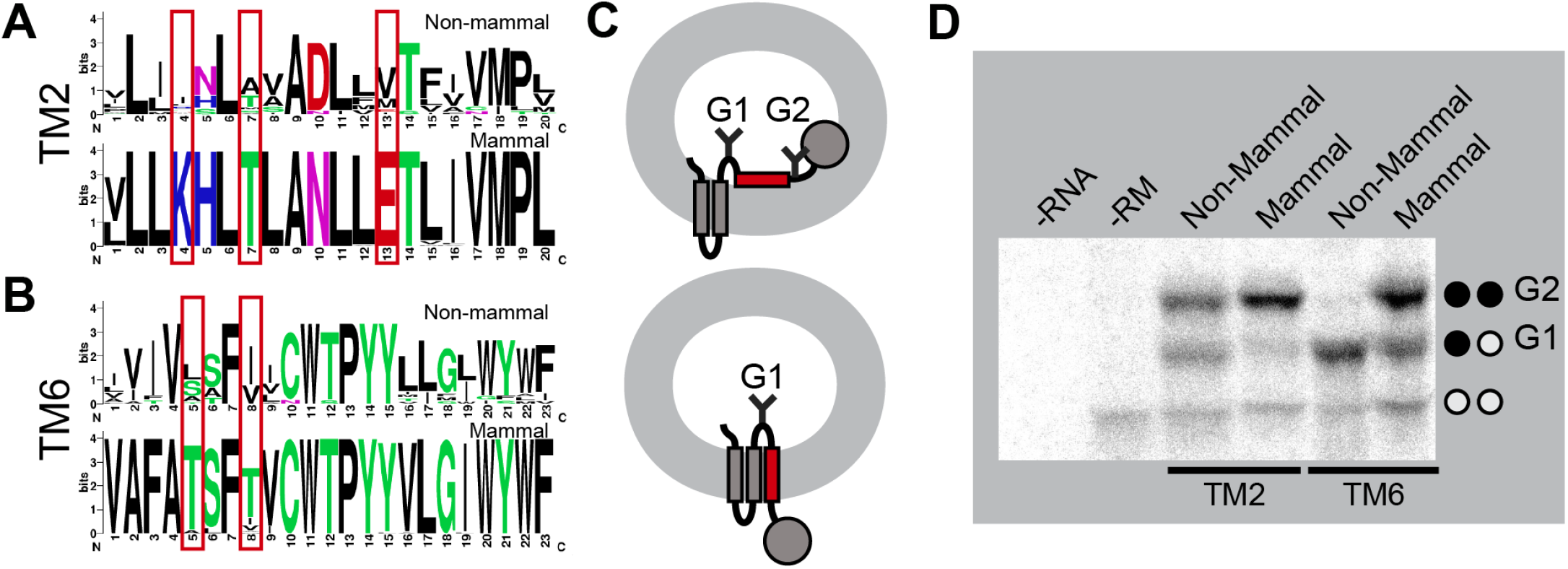
Translocon-mediated membrane integration of TMs 2 & 6. Differences in the sequences of TM2 and TM6 are analyzed in relation to differences in their efficiency of translocon-mediated membrane integration. A) Logo plots depict the most common amino acid at each position within TM2 of non-mammalian (top) and mammalian (bottom) GnRHRs. The positions of polar substitutions are indicated with a red box. B) Logo plots depict the most common amino acid at each position within TM6 of non-mammalian (top) and mammalian (bottom) GnRHRs. The positions of polar substitutions are indicated with a red box. C) A cartoon depicts the manner in which the translocon-mediated membrane integration of the guested TM domain within chimeric Lep proteins impacts their glycosylation. A failure of the guest TM domain to undergo translocon-mediated membrane integration results in the glycosylation of two residues (top), whereas the membrane integration of the guest domain results in a single glycosylation (bottom). D) Chimeric Lep proteins containing the mammalian or non-mammalian consensus sequences for TM2 and TM6 were translated in canine rough microsomes and analyzed by SDS-PAGE. Negative control reactions lacking RNA (first lane) or containing RNA but lacking microsomes (second lane) are shown for reference. The positions of the untargeted (no glycans), singly glycosylated (G1), and doubly glycosylated (G2) forms of the protein are indicated for reference.

### Structural Context of Polar Residues and Their Impacts on PME

Logo plots show that several polar residues were introduced within TM2 and TM6 during the evolutionary adaptation of mammalian GnRHRs (Figure 4 A & B). Though these mutations disrupt cotranslational folding (Figure 4D), it is possible that they also help to stabilize the structure of the folded receptor and/or enhance its function. To gain insights into the structural context of these residues, we constructed comparative models of both the cGnRHR and hGnRHR receptors. With the exception of the C-terminal tail of cGnRHR, both forms of the receptor have a similar architecture (Figure 5A, Cα RMSD 2.95 Å). The model of hGnRHR reveals that each of the three polar substitutions within TM2 occur at position that are buried within the protein core (Figure 5B). Thus, these side chains are likely to play integral roles in the native structure and/ or function of mammalian GnRHRs. In contrast, the two polar substitutions within TM6 appear to have occurred at surface residues that are exposed to the hydrophobic core of the lipid bilayer (Figure 5C). Though a complex role of these side chains in the native conformational dynamics cannot be ruled out, this observation raises the possibility that these mutations primarily tune the fidelity of cotranslational folding and the corresponding PME without impacting function.

**Figure 5.**
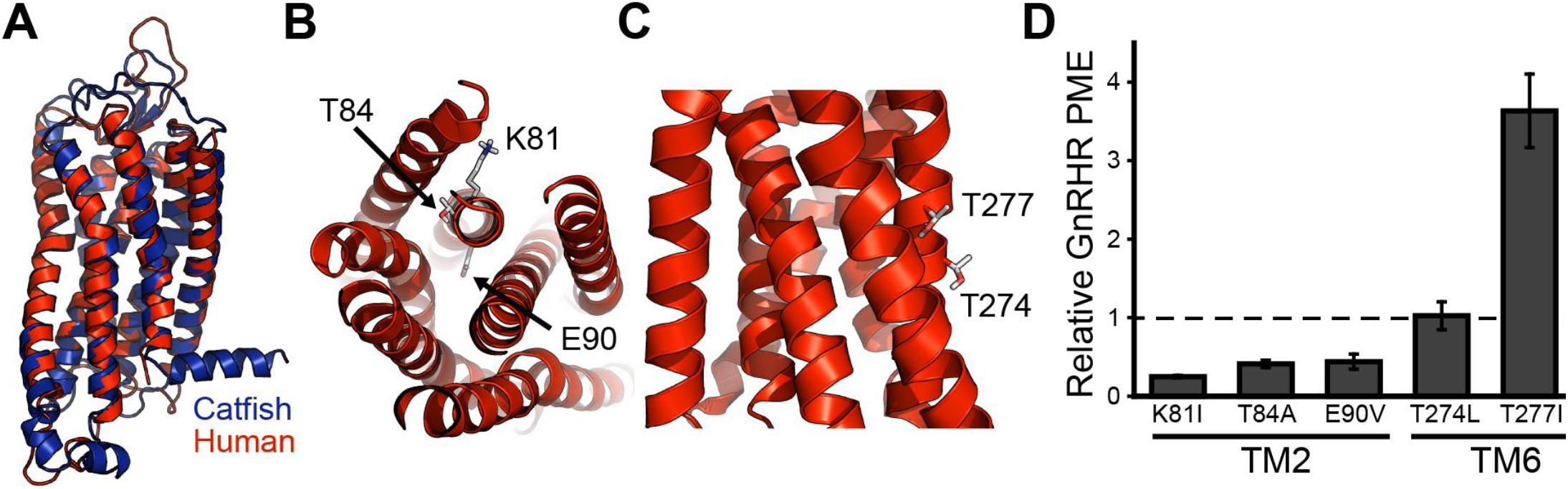
Structural context and proteostatic effects of polar residues within TMs 2 & 7. Structural models of catfish (blue) and human (red) GnRHR illustrate the structural context of polar side chains within TMs 2 & 6, and the effects of hydrophobic substitutions at these positions on the plasma membrane expression (PME) of human GnRHR is shown. A) Structural homology models of human (red) and catfish (blue) GnRHRs are overlaid for reference. B) A cutaway of the shows that the polar residues of interest within TM2 (K81, T84, and E90) are buried within the core of the human GnRHR protein. C) A side view shows that the polar residues of interest within TM6 of human GnRHR appear to be projected into the membrane core. D) Polar side chains within TMs 2 & 6 were replaced with the most common hydrophobic residues found within non-mammalian GnRHRs, and the effects of these substitutions one the plasma membrane expression (PME) of human GnRHR was measured in HEK293T cells by flow cytometry. A bar graph depicts the mean fluorescence intensity associated with the surface immunostaining of each variant normalized relative to that of WT human GnRHR. Values reflect the average of three biological replicates, and error bars reflect the standard deviation.

If the inefficient cotranslational membrane integration of TM2 and/or TM6 contributes to the proteostatic adaptations in mammalian GnRHRs, then substitutions that enhance the hydrophobicity of these domains could potentially restore their PME. We therefore measured the PME of a series of hGnRHR variants containing ancestral substitutions that partially restore the hydrophobicity of TM2 and TM6. In each case, replacing polar residues in TM2 with the hydrophobic consensus non-mammalian residue resulted in a further reduction in the PME of hGnRHR (Figure 5D). This result is perhaps unsurprising. While these substitutions are likely to enhance cotranslational folding, the structural model of hGnRHR suggests they are also likely to introduce packing defects in the native structure (Figure 5B). Thus, the structural basis for the net proteostatic impact of these substitutions is unclear. In contrast, consensus hydrophobic substitutions at surface residues within TM6 appear to be well-tolerated. Though T274L has no impact on PME, replacing T277 with an isoleucine enhances the PME of GnRHR 3.6 ± 0.5 fold (Figure 5D). Thus, restoring a hydrophobic side chain to a single lipid-exposed residue within TM6 (Figure 5C) is sufficient to partially recover the PME of hGnRHR. In conjunction with *in vitro* translation measurements (Figure 4D), this observation implies that the enhanced cotranslational misfolding of mammalian TM6 contributes to the attenuated PME of these proteins.

### Functional Impact of the T277I Substitution

Our results collectively reveal that the enhanced polarity of TM6 compromises the cotranslational folding and expression of mammalian GnRHRs, and that the reversion of a single surface residue to its ancestral hydrophobic side chain (T277I) is sufficient to partially recover PME. Nevertheless, it is possible that this side chain was introduced to support GnRHR function. To determine whether this residue is important for GnRHR signaling, we compared the activity of WT and T277I hGnRHRs. Briefly, cells transiently expressing WT or T277I GnRHR were titrated with gonadotropin-releasing hormone (GnRH), and the receptor activation was indirectly measured by the magnitude of the resulting cytosolic calcium flux. Cells expressing these receptors exhibit robust response to GnRH, and the magnitude of the calcium flux was quite similar for WT and T277I hGnRHR (Figure 6A). Moreover, the fitted EC_50_ values for WT (0.61 ± 0.38 μM) and T277I (0.23 ± 0.18 μM) were found to be statistically indistinguishable (Figure 6A). These findings demonstrate that T277 is not essential for hGnRHR function. Given that this non-essential polar side chain negatively impacts cotranslational folding and PME, our collective observations suggest that the evolved polarity of this segment serves to tune the PME of mammalian GnRHRs.

**Figure 6.**
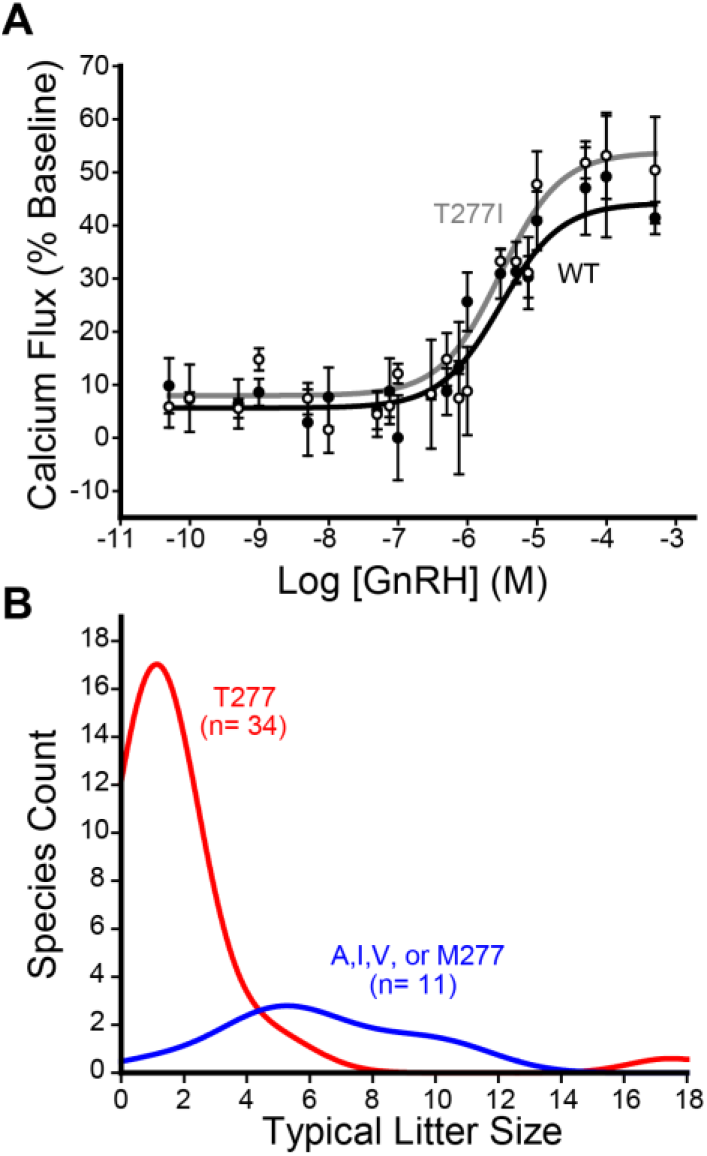
Hydrophobicity of Residue 277 in Relation to GnRHR Function and Mammalian Litter Size. A) The activation of WT and T277I hGnRHR was measured in response to varying doses of human gonadotropin releasing hormone (GnRH) in HEK293T cells. Cells transiently expressing each receptor were stimulated with GnRH, and signaling was measured by the change in the fluorescence intensity of a cytosolic calcium reporter. The average fluorescence intensities of cells expressing WT (●) or T277I (○) from three technical replicates are normalized relative to baseline and plotted against the corresponding concentration of hormone. Error bars reflect the standard deviation from three technical replicates. Curves reflect the fit of the WT (black) and T277I (gray) data to a single-site binding model. B) A histogram depicts the distribution of litter sizes for mammals bearing a polar (red) or hydrophobic (blue) side chain at residue 277.

### Sequence Variations in Relation to Reproductive Outcomes

Evolutionary variations in PME of GnRHR should have a direct influence on GnRH signaling, which may alter reproductive outcomes. Our results suggest that the hydrophobicity of residue 277 modulates the PME of GnRHR. To determine whether natural variation at this position coincides with differences in reproductive outcomes, we compared the hydrophobicity of the side chain at this position to litter size among 44 mammalian species. Species with a hydrophobic residue at this position have significantly larger litters than those with a hydrophilic residue (Figure 6B, Mann-Whitney *p* = 1.7 × 10^−5^), suggesting that the polarity of TM6 is associated with reproductive traits at the organismal level. Moreover, of the polar residues within TM2 and TM6, only residue 277 exhibits appreciable variation in hydrophobicity among the mammalian forms of the receptor (Figure 4 A & B). These observations potentially suggest that modifications at T277, and their effects on the PME of GnRHR, may have played a direct role in the optimization of reproductive fitness.

## Discussion

Previous investigations on the evolution of GnRHR have suggested that its activity has been down-regulated in mammals through various coding mutations that enhanced its propensity to misfold (*5*, *14*, *16*). In this work, we followed up on these investigations in order to assess the molecular basis for this evolved instability. Using quantitative cellular measurements, we first confirmed that the catfish receptor exhibits more robust expression and cellular trafficking relative to mammalian GnRHRs. To determine which mutations contributed to the lapse in mammalian GnRHR proteostasis, we then characterized the effects of various sequence modifications on the PME of cGnRHR. Our results provide additional evidence that the C-terminal truncation in mammalian GnRHRs is likely a key factor leading to their attenuated PME (Figure S1). However, an analysis of known GnRHR sequences also shows that the TM domains of mammalian GnRHRs are more polar than their non-mammalian counterparts (Figures 2 & 3). We find that this increased polarity compromises the translocon-mediated cotranslational folding of at least two TM domains (Figure 4). Moreover, we show that restoring a hydrophobic amino acid at a surface residue within TM6 increases the PME of hGnRHR 3.6-fold with no impact on receptor activation (Figures 5D & 6A). This modification is likely relevant to the evolutionary trajectory of GnRHR considering the mouse receptor, which exhibits a higher PME than the human receptor, features a hydrophobic residue at this position (V276). Indeed, the hydrophobicity of this residue is associated with differences in mammalian litter sizes (Figure 6B).

There are several caveats to these investigations. First, it should be noted that epistatic interactions between some of these mutations may alter their effects on PME in the context of mammalian receptors. Furthermore, we cannot measure the PME of these receptors in the context of their native environment within the pituitary gland of each animal. It is likely that the magnitude of these proteostatic effects are distinct in the context of the native proteostasis networks that typically support GnRHR biogenesis. Nevertheless, to our knowledge mammalian GnRHRs are the only class A GPCRs that completely lack helix 8 and/ or a C-terminal tail (*18*). We suspect this drastic modification is likely to have consequences for the maturation and sorting of mammalian GnRHRs within the secretory pathway regardless of the cellular context. It should also be noted that the mechanism of the translocon is highly conserved, and the increased polarity of TM2 and TM6 are therefore likely to reduce the efficiency of cotranslational GnRHR folding in any cellular context (*23*). Indeed, the hydrophobicity of TM domains is also known to be a critical factor that governs the expression of membrane proteins in *E. coli* (*24*, *25*). Based on these considerations, it seems likely that both the C-terminal truncation and the enhanced the polarity of the TM domains of GnRHR are likely to have contributed to evolutionary modifications to the PME and net activity of mammalian GnRHRs.

Our findings provide additional evidence to suggest the activity of mammalian GnRHRs has been tuned through modulation of GnRHR folding rather than through transcriptional modifications. This is perhaps surprising given the metabolic cost of protein synthesis. Nevertheless, we believe this outcome is reasonable in light of certain evolutionary considerations. It should first be noted that the length of the open reading frame of GnRHR is roughly six times that of its promoter (*26*). Considering most coding variants are destabilizing, there are likely to be far more coding variants that decrease the PME of the receptor relative to the number that would simply decrease its transcription. If selection pressures simply favored attenuated GnRHR signaling, then it is perhaps most probable this would arise from mutations that destabilize the native GnRHR structure. Consistent with the observed proteostatic patterns (Figure 1), such mutations would result in both a decreased PME and an increased accumulation of the receptor within the secretory pathway. Though energetically wasteful, it is unclear whether the biosynthesis of misfolded GnRHRs necessarily imposes a significant fitness burden in this case, as this receptor is only expressed at moderate levels within the pituitary gland according to the human protein atlas (proteinatlas.org). Thus, it seems plausible that various destabilizing mutations fixed in mammals provided the gain in reproductive fitness outweighs the metabolic costs associated with the synthesis and degradation of misfolded receptors.

Marginal conformational stability is an emergent property of naturally evolved proteins (*2*, *27*, *28*). This instability has been previously attributed to the net-destabilizing effects of random mutations in conjunction with a general lack of selection pressure for hyper-stable proteins (*2*). Although it is likely that the instability of GnRHR emerged as a result genetic drift, the net variation in PME resulting from these mutations was likely constrained by adaptive changes in reproductive fitness. Mammals, which have fewer offspring at higher metabolic cost, may require less GnRHR activity than non-mammals to maintain reproduction. The loss of the C-terminal tail and the increased polarity of the TM domains, while deleterious to folding and PME, may have been tolerated due to an attenuated reliance on GnRH signaling. Nevertheless, it is certainly possible that the incorporation of polar residues into TM2 and TM6, and the resulting variation in GnRHR PME, played an active role in the optimization of mammalian reproductive fitness. In the context of integral MPs, the evolutionary utility of sub-optimal cotranslational folding energetics is illustrated by the fact that the PME of rhodopsin is highly sensitive a wide variety of mutations within its TM domains (*29*). The natural exploitation of the thin energetic margins involved in the cotranslational MP folding perhaps also provides an explanation for the fact that the hydrophobicity of rhodopsin’s TM domains has not been optimized to promote efficient biosynthesis (*4*). However, the apparent malleability of cotranslational MP folding energetics does not come without costs, as TM domains are generally less tolerant of genetic variation and mutations within TM domains give rise to numerous genetic diseases (*3*, *30*, *31*). Together, these observations provide new insights into the molecular mechanisms of membrane protein evolution.

## Materials & Methods

### Plasmid Preparation and Mutagenesis

A series of pcDNA5 FRT expression vector containing various GnRHR cDNAs containing an N-terminal influenza hemaglutinin (HA) epitope were used for the transient expression of GnRHR variants. GnRHR cDNAs in this vector are followed by an internal ribosome entry site (IRES) and eGFP sequence, which generates bicistronic GFP expression in positively transfected cells. Vectors containing GnRHRs from various species were generated using In-Fusion HD Cloning (Takara Bio, Shiga, Japan). Mutations were introduced by site-directed mutagenesis, and truncations were generated by In-Fusion HD cloning. To adapt these constructs for functional experiments, the HA epitope was deleted to minimize interferences with ligand binding, and the IRES eGFP sequence was deleted to prevent interference of eGFP with fluorescence measurements.

A previously described pGEM expression vector containing modified leader peptidase (Lep) cDNA was used for *in vitro* translation (*22*). TM domains of interest were cloned into the H-segment position within the Lep gene using directional cloning at the *Spe*I and *Kpn*I restriction sites. All plasmids were prepared with the Endotoxin-Free Zymopure Midiprep or Miniprep Kit (Zymo Research, Irvine, CA).

### In vitro Translation of Chimeric Lep Proteins

Messenger RNA (mRNA) was generated for each chimeric Lep gene using the RiboMAX SP6 kit (Promega, Madison, WI). Lep variants were then translated using rabbit reticulocyte lysate (Promega, Madison, WI) supplemented with canine rough microsomes (tRNA probes, College Station, TX), and EasyTag ^35^S-labeled methionine (PerkinElmer, Waltham, MA). Translation was carried out at 30 ^o^C for 60 minutes. Reactions were diluted 1:4 in 1X SDS PAGE loading buffer and separated on a 12% SDS PAGE gel. Gels were then dried, exposed overnight on a phosphor imaging plate (GE Healthcare, New York, NY), and imaged on a Typhoon Imager (GE Healthcare, New York, NY). The ratio of singly (*G1*) to doubly (*G2*) glycosylated Lep protein was quantified by densitometry using ImageJ software. The *G1:G2* ratio represents an apparent equilibrium constant (*K_app_*) for the transfer of the H-segment from the translocon to the membrane, as previously described. Apparent transfer free energy values for the H-segments were calculated using the following equation:

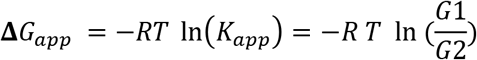

where Δ*G*_*app*_ represents the apparent free energy for the transfer of the H-segment into the membrane, *R* represents the universal gas constant, *T* represents the temperature, *K*_*app*_ represents the apparent equilibrium constant for the transfer of the H-segment from the translocon into the membrane, *G1* represents the intensity of the singly glycosylated band and *G2* represents the intensity of the doubly glycosylated band, as was previously described (*21*). Reported transfer free energy values represent the average values from three experimental replicates.

### Cellular GnRHR Expression Measurements

To quantitatively measure the cellular trafficking of GnRHR variants, plasma membrane and intracellular GnRHRs were differentially immunostained and analyzed by flow cytometry, as described previously (*17*). GnRHR variants were transiently expressed in HEK293T cells using Lipofectamine 3000 (Invitrogen, Carlsbad, CA). Two days after transfection, the cells were harvested with TrypLE Express protease (Gibco, Grand Island, NY). Plasma membrane GnRHRs were then immunostained for 30 minutes in the dark with a DyLight 550-conjugated anti-HA antibody (Invitrogen, Carlsbad, CA). Cells were fixed and permeabilized using a Fix and Perm kit (Invitrogen, Carlsbad, CA), and washed twice with 2% fetal bovine serum in phosphate-buffered saline (wash buffer). Intracellular GnRHRs were then immunostained for 30 minutes in the dark using an Alexa Fluor 647-conjugated anti-HA antibody (Invitrogen, Carlsbad, CA). Cells were washed twice in order to remove excess antibody prior to analysis of cellular fluorescence profiles. Fluorescence profiles were analyzed on a BD LSRII flow cytometer (BD Biosciences, Franklin Lakes, NJ). Forward and side scatter measurements were used to set a gate for intact single cells. eGFP intensities (488 nm laser, 530/30 nm emission filter) were then used to set a gate for positively-transfected cells. DyLight 550 (561 nm laser, 582/15 nm emission filter) and Alexa Fluor 647 (640 nm laser, 670/30 nm emission filter) intensities were then calculated for several thousand positively-transfected single cells within each biological replicate. Data were analyzed using FlowJo software (Treestar, Ashland, OR). Characterizations of GnRHR variant expression levels were carried out with at least three biological replicates each.

### Functional Measurements of GnRHRs

GnRHR activity was measured in HEK293T cells by monitoring cytosolic calcium fluxes that occurred in response to gonadotropin-releasing hormone (GnRH, Sigma Aldrich, St. Louis, MO). GnRHR variants were transiently expressed in HEK293T cells using Lipofectamine 3000 (Invitrogen, Carlsbad, CA). Two days after transfection, the cells were harvested with TrypLE Express (Gibco, Grand Island, NY), then re-plated in 96-well plates (Corning, Big Flats, NY) coated with poly-D-lysine (Gibco, Grand Island, NY) at a density of 60,000 cells per well. Cells were then dosed the following day and assayed using the FLIPR Calcium 6-QF Assay Kit (Molecular Devices, San Jose, CA) according to the manufacturer’s protocol. The fluorescence intensities of cells incubated in the calcium-sensitive FLIPR dye was measured for thirty seconds prior to dosing with GnRH using a Synergy Neo2 microplate reader (BioTek, Winooski, VT) using an excitation wavelength of 485/20 nm and an emission filter at 525/10 nm. Directly after dosing the cells, the change in fluorescence was measured for six minutes. The percent calcium flux under each condition was calculated for each well using the following equation:

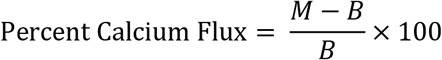

where *M* is the maximum fluorescence value for the calcium flux and *B* is the baseline signal as was determined from by averaging the fluorescence intensity before ligand addition. EC_50_ values were determined by fitting titrations to the following function:

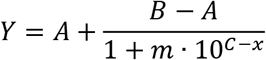

where *Y* is the percent calcium flux, *A* is the minimal curve asymptote, *B* is the maximal curve asymptote, *m* is the slope of the transition region, *C* is the logarithm of the EC_50_, and *x* is the logarithm of the GnRH concentration (*32*). Reported EC_50_ values represent the average from three biological replicates.

### Selection and Analysis of GnRHR Sequences

59 GnRHR sequences from different species were collected from the NCBI (https://www.ncbi.nlm.nih.gov) and Uniprot (https://www.uniprot.org) databases. Humans have only one type of GnRHR (GnRHR-I), while other species may have multiple types (*33*, *34*). Sequences selected for phylogenetic analysis were therefore limited to those annotated as GnRHR-I in order to analyze trends across species.

A phylogenetic tree was generated from these sequences using MEGA7 software (megasoftware.net). A sequence alignment was generated using the MUSCLE alignment tool with default settings. This alignment was then used to construct a Maximum Likelihood tree (*35*). The positions of nascent TM domains within each sequence were then identified from energetic minima generated with a window scan function within the ΔG predictor, which sums depth-dependent free energies associated with the transfer of amino acids from the translocon to the ER membrane (http://dgpred.cbr.su.se/) (*22*). The ΔG predictor was then used to calculate the free energy difference associated with the translocon-mediated membrane integration of each putative TM domain (*22*). The phylogenetic tree and ΔG prediction data were then uploaded to the Interactive Tree of Life (https://itol.embl.de), where the ΔG predictions were displayed as color gradients on the phylogenetic tree (*36*).

To generate logo plots, the GnRHR-I sequences were first aligned in ClustalOmega (*37*). The positions of the TM domains within the hGnRHR sequence were determined by the ΔG predictor, and the transmembrane domains in other species were then identified by the corresponding positions in the alignment. Sequence logo plots were then generated for each transmembrane domain in the mammalian and non-mammalian sequences using the WebLogo application (https://weblogo.berkeley.edu/logo.cgi) (*38*).

Litter size data was collected for 44 species corresponding to the mammalian GnRHR-I sequences. The average litter size or the middle of a range was used as the typical litter size. For species where twins or multiples are rare, the typical litter size was set to one. The residue equivalent to T277 in hGnRHR was determined for each mammalian GnRHR-I sequence by an alignment in ClustalOmega (*37*).

### Structural Modeling

Comparative models of the human and catfish forms GnRHR were generated using multi-template comparative modeling in Rosetta. The GnRHR sequence was first aligned with sequences for 34 GPCR crystal structures obtained from GPCRdb (http://www.gpcrdb.org) (*39*). Manual adjustments were then made to account for well-known conserved residues in loop regions and TM domains (*40*). OCTOPUS was used to define the TM domains, and the two disulfide bonds were defined manually (*41*, *42*). To generate a model of GnRHR in the inactive state, the sequences were threaded onto the antagonist-bound structures of several other Class A Group β GPCRs including the human OX2 orexin receptor (HCRTR2, PDB 4S0V, 2.5 Å), human OX1 orexin receptor (HCRTR1, PDB 4ZJC, 2.8 Å), human endothelin receptor type B (EDNRB, PDB 5×93, 2.2 Å), and human neuropeptide Y receptor Y1 (NPY1R, PDB 5ZBQ, 2.7 Å). Threading was completed using the *partial thread* application in RosettaCM (*43*). 1,000 models were then generated using the *hybridize* application in RosettaCM and the TM domains were relaxed using a set of optimized RosettaMembrane weights that were modified from the Talaris scoring function (*44*). The models with the lowest Rosetta energy score were used for structural analysis.

## Acknowledgments

We thank Christiane Hassel and the Indiana University Bloomington Flow Cytometry Core Facility for their experimental support. This research was supported in part by grant from the National Institute of General Medical Sciences (NIGMS) under award numbers R01GM129261, R01GM080403, R01HL122010, and R01DA046138. This work was also supported by a National Science Foundation (NSF) Graduate Research Fellowship to L.M. C. (grant number 1342962). Any opinions, findings, and conclusions or recommendations expressed in this material are those of the author(s) and do not necessarily reflect the views of the NSF or NIH.

