## Supplemental Materials for "Molecular Basis for the Evolved Instability of a Human G-Protein Coupled Receptor"

This document includes:
-Figure S1
-Figure S2
-Table S1
-Table S2
-Table S3
-Supplemental References

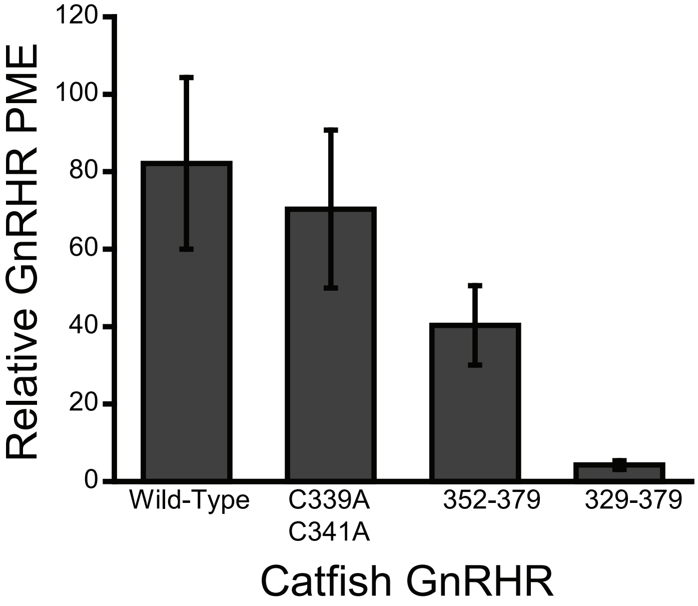


**Figure S1. Impact of C-terminal modifications on the plasma membrane expression of cGnRHR.** Catfish GnRHRs bearing C-terminal modifications were transiently expressed in HEK293T cells prior to analysis of surface immunostaining of plasma membrane GnRHR by flow cytometry. A bar graph depicts the mean fluorescence intensity associated with the surface immunostaining of a series of catfish GnRHR variants normalized relative to that of human GnRHR. Values reflect the average of three biological replicates, and error bars reflect the standard deviation.

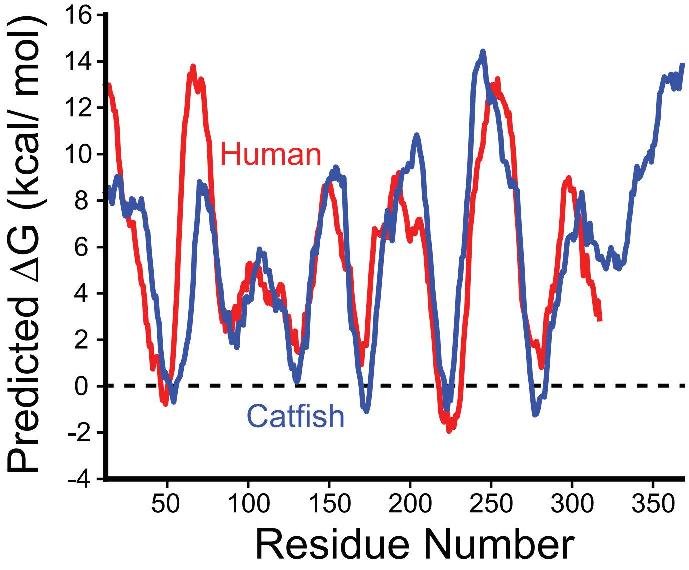


**Figure S2. Topological energetics of human and catfish GnRHRs.** Sequence-based predictions of the topological energetics of human GnRHR (red) are analyzed in relation to those of catfish GnRHR (blue). The predicted transfer free energy from the translocon to the ER membrane was predicted for each possible 23-residue segment within each protein were calculated using the ΔG predictor, and each predicted value was plotted as a function of the central residue of the segment. A ΔG value of 0 kcal/ mol, at which membrane integration is energetically neutral, is indicated with a dashed line.

**Table S1. Predicted transfer free energies associated with the putative TM domains within GnRHRs.**

| | | Predicted Transfer Free Energy (kcal/mol) | | | | | | | |
|---|---|---|---|---|---|---|---|---|---|
| **Species** | **Accession ID** | **TM1** | **TM2** | **TM3** | **TM4** | **TM5** | **TM6** | **TM7** | **7 TM Average** |
| *Sebastes schlegelii* | AFR51715.1 | -0.291 | 0.825 | 0.906 | 1.105 | -2.699 | -0.554 | 0.876 | 0.024 |
| *Octopus vulgaris* | Q2V2K5.1 | -0.028 | 1.189 | 1.424 | 0.542 | -2.913 | -1.049 | 2.172 | 0.191 |
| *Xenopus tropicalis* | NP_001107549.1 | -0.637 | -0.343 | 1.274 | 0.538 | -3.978 | -1.698 | 1.694 | -0.450 |
| *Callorhinchus milii* | NP_001279833.1 | -1.615 | 2.202 | 1.382 | -1.329 | -2.228 | -0.411 | 3.168 | 0.167 |
| *Dicentrarchus labrax* | CAD11992.1 | -0.816 | 0.825 | 0.906 | 0.663 | -2.699 | -0.323 | 1.166 | -0.040 |
| *Danio rerio* | XP_021323652.1 | -0.621 | 1.556 | -0.264 | -0.995 | -1.031 | -1.108 | 4.667 | 0.315 |
| *Macrobrachium nipponense* | AHB33640.1 | 0.957 | 0.78 | 2.604 | -0.513 | 0.652 | -2.433 | -2.182 | -0.019 |
| *Acanthopagrus schlegelii* | AAV71128.1 | -0.481 | 0.825 | 0.906 | 0.895 | -3.319 | -0.554 | 1.166 | -0.080 |
| *Branchiostoma floridae* | ACC68665.1 | -0.883 | 0.546 | -1.065 | 2.711 | -3.374 | -0.159 | 3.467 | 0.178 |
| *Odontesthes bonariensis* | ABI75337.1 | -1.482 | 1.45 | -0.229 | -1.171 | -0.825 | -1.1 | 4.821 | 0.209 |
| *Kryptolebias marmoratus* | ABK88281.1 | -0.311 | 0.826 | -0.1 | 0.505 | -2.422 | -1.499 | 1.021 | -0.283 |
| *Python bivittatus* | XP_007437330.1 | 1.195 | 0.044 | 0.089 | 0.121 | -3.593 | -0.133 | 1.454 | -0.118 |
| *Monopterus albus* | ARS88253.1 | -1.077 | 0.825 | 0.906 | 1.418 | -2.699 | -1.39 | 2.867 | 0.121 |
| *Cairina moschata* | AGO01050.1 | 0.032 | 1.003 | -0.425 | -0.177 | -3.323 | -0.72 | 1.337 | -0.325 |
| *Gallus gallus* | AID62089.1 | 0.01 | 0.897 | -0.425 | 0.082 | -2.922 | -0.634 | 1.785 | -0.172 |
| *Ovis aries* | NP_001009397.1 | -0.854 | 2.352 | 1.334 | 0.677 | -2.053 | 0.805 | 3.015 | 0.754 |
| *Bos Taurus* | NP_803480.1 | -0.854 | 2.165 | 1.334 | 0.677 | -1.754 | 0.805 | 2.566 | 0.706 |
| *Canis lupus familiaris* | NP_001003121.1 | -1.053 | 2.352 | 1.334 | 0.305 | -1.523 | 0.805 | 2.419 | 0.663 |
| *Bos mutus grunniens* | Q19PY9.1 | -0.854 | 2.165 | 1.334 | 0.677 | -1.754 | 0.805 | 2.117 | 0.641 |
| *Sus scrofa* | NP_999438.1 | -0.646 | 2.352 | 1.334 | 0.341 | -1.949 | -0.748 | 2.566 | 0.464 |
| *Cavia porcellus* | NP_001166428.1 | -0.363 | 2.352 | 1.643 | 2.21 | -2.785 | -0.163 | 2.566 | 0.780 |
| *Trichosurus vulpecula* | Q9TTI8.1 | -0.267 | 2.456 | 0.655 | -0.773 | -1.129 | -0.159 | 2.87 | 0.522 |
| *Delphinapterus leucas* | XP_022417405.1 | -0.38 | 2.352 | 1.179 | 0.306 | -2.389 | 0.805 | 2.566 | 0.634 |
| *Ictidomys tridecemlineatus* | XP_005331769.1 | -0.188 | 2.352 | 1.334 | 0.942 | -2.053 | 0.046 | 2.566 | 0.714 |
| *Neomonachus schauinslandi* | XP_021558093.1 | -0.38 | 2.497 | 1.334 | 0.305 | -1.624 | 0.563 | 2.419 | 0.731 |
| *Phascolarctos cinereus* | XP_020855496.1 | -0.267 | 2.456 | 0.655 | -0.134 | -1.784 | 0.08 | 3.176 | 0.597 |
| *Odocoileus virginianus texanus* | XP_020769734.1 | -0.854 | 2.352 | 1.334 | 0.677 | -1.754 | 0.805 | 2.566 | 0.732 |
| *Bubalus bubalis* | ARJ58634.1 | -0.854 | 2.165 | 1.334 | 0.677 | -1.754 | 0.805 | 2.566 | 0.706 |
| *Capra hircus* | NP_001272541.1 | -0.854 | 2.352 | 1.334 | 0.677 | -2.053 | 0.805 | 2.566 | 0.690 |
| *Equus caballus* | NP_001075305.1 | -0.38 | 2.352 | 1.334 | 0.535 | -1.884 | 0.805 | 2.419 | 0.740 |
| *Felis catus* | AFP97799.1 | -0.38 | 2.316 | 1.334 | 0.389 | -2.053 | 0.805 | 2.419 | 0.690 |
| *Sarcophilus harrisii* | XP_003773267.1 | -0.477 | 2.475 | 0.655 | 0.101 | -1.947 | 0.08 | 2.87 | 0.537 |
| *Loxodonta Africana* | XP_003415950.1 | -0.767 | 2.165 | 1.832 | 0.764 | -1.744 | 0.805 | 2.566 | 0.803 |
| *Dasypus novemcinctus* | XP_004466930.1 | -0.249 | 2.165 | 1.334 | 0.787 | -1.901 | 0.303 | 2.566 | 0.715 |
| *Pteropus vampyrus* | XP_011358956.1 | -0.293 | 2.352 | 1.334 | 0.773 | -1.481 | 0.805 | 2.566 | 0.865 |
| *Myotis lucifugus* | XP_006095704.1 | -0.636 | 2.352 | 1.334 | 0.639 | -1.837 | 0.805 | 2.566 | 0.746 |
| *Physeter catodon* | XP_007108070.1 | -0.38 | 2.352 | 1.179 | 0.306 | -2.504 | 0.805 | 2.566 | 0.618 |
| *Desmodus rotundus* | XP_024433366.1 | -0.72 | 2.352 | 1.643 | 0.251 | -1.837 | 0.563 | 2.566 | 0.688 |
| *Neophocaena asiaeorientalis* | XP_024603511.1 | -0.38 | 2.352 | 1.179 | 0.306 | -2.389 | 0.805 | 2.566 | 0.634 |
| *Pteropus alecto* | XP_006916576.1 | -0.38 | 2.352 | 1.334 | 0.773 | -1.481 | 0.805 | 2.566 | 0.853 |
| *Canis lupus dingo* | XP_025291626.1 | -1.053 | 2.352 | 1.334 | 0.305 | -1.523 | 0.805 | 2.419 | 0.663 |
| *Oryctolagus cuniculus* | AAV48838.1 | -0.091 | 2.352 | 1.334 | 0.431 | -2.102 | 0.046 | 2.566 | 0.648 |
| *Rattus norvegicus* | NP_112300.2 | -0.361 | 2.352 | 1.461 | -0.262 | -2.053 | 0.037 | 2.566 | 0.534 |
| *Mus musculus* | Q01776.1 | -0.646 | 2.352 | 1.334 | -0.262 | -2.18 | 0.26 | 2.566 | 0.489 |
| *Heterocephalus glaber* | XP_004849709.1 | -0.401 | 2.352 | 1.03 | 1.501 | -2.053 | 0.046 | 2.566 | 0.720 |
| *Mus caroli* | XP_021019370.1 | -0.646 | 2.352 | 1.334 | -0.262 | -2.18 | 0.26 | 2.566 | 0.489 |
| *Mus pahari* | XP_021066838.1 | -0.646 | 2.352 | 1.334 | -0.29 | -2.18 | 0.26 | 2.566 | 0.485 |
| *'Papio anubis* | XP_003898885.1 | -0.774 | 2.165 | 1.334 | 0.934 | -2.053 | 0.805 | 2.566 | 0.711 |
| *Carlito syrichta* | XP_008069339.1 | -1.126 | 2.352 | 2.049 | 0.378 | -2.053 | 0.805 | 2.419 | 0.689 |
| *Aotus nancymaae* | XP_012328857.1 | -0.38 | 2.165 | 0.655 | 0.934 | -2.053 | 0.805 | 2.566 | 0.670 |
| *Microcebus murinus* | XP_012616420.1 | -0.453 | 2.352 | 1.334 | 0.764 | -2.376 | 0.805 | 2.566 | 0.713 |
| *Theropithecus gelada* | XP_025242022.1 | -0.845 | 2.165 | 1.239 | 0.934 | -2.053 | 0.805 | 2.566 | 0.687 |
| *Pan paniscus* | XP_003815858.1 | -0.774 | 2.165 | 1.334 | 0.934 | -1.943 | 0.805 | 2.566 | 0.727 |
| *Macaca nemestrina* | XP_011709114.1 | -0.884 | 2.165 | 1.334 | 0.934 | -2.053 | 0.805 | 2.566 | 0.695 |
| *Pan troglodytes* | XP_526608.1 | -0.774 | 2.165 | 1.334 | 0.934 | -1.943 | 0.805 | 2.566 | 0.727 |
| *Pongo abelii* | XP_024101999.1 | -0.774 | 2.165 | 1.334 | 0.934 | -2.053 | 0.805 | 2.566 | 0.711 |
| *Piliocolobus tephrosceles* | XP_023087932.1 | -0.774 | 2.165 | 1.334 | 0.934 | -2.053 | 0.805 | 2.566 | 0.711 |
| *Otolemur garnettii* | XP_003801017.1 | -0.38 | 2.336 | 2.105 | 0.856 | -2.247 | 0.673 | 2.566 | 0.844 |
| *Homo sapiens* | AAB26287.1 | -0.774 | 2.165 | 1.334 | 0.934 | -1.943 | 0.805 | 2.566 | 0.727 |

**Table S2. Predicted versus measured apparent transfer free energies associated with TMs 2 & 6.**

| TM Segment | Predicted $\Delta G_{app}$ (kcal/ mol)† | Measured $\Delta G_{app}$ (kcal/ mol)‡ |
|---|---|---|
| Non-mammalian TM2 | 0.73 | 0.15 ± 0.08 |
| Mammalian TM2 | 2.42 | 0.85 ± 0.1 |
| Non-mammalian TM6 | -0.26 | -1.38 ± 0.01 |
| Mammalian TM6 | 0.81 | 0.23 ± 0.05 |

†Values were predicted from sequence using the ΔG Predictor.
‡Values reflect the average from three independent experimental replicates, and errors reflect the standard deviation.

### Table S3. Litter size data associated with mammalian GnRHRs.

| Species | Accession ID | Residue at Position 277 | Typical Litter Size |
|---|---|---|---|
| *Ovis aries* | NP_001009397.1 | T | 1.58 (*1*) |
| *Bos Taurus* | NP_803480.1 | T | 1 (*1*) |
| *Canis lupus familiaris* | NP_001003121.1 | T | 5.4 (*2*) |
| *Bos mutus grunniens* | Q19PY9.1 | T | 1 (*3*) |
| *Sus scrofa* | NP_999438.1 | I | 5 (*4*) |
| *Cavia porcellus* | NP_001166428.1 | I | 3.8 (*1*) |
| *Trichosurus vulpecula* | Q9TTI8.1 | T | 1 (*5*) |
| *Delphinapterus leucas* | XP_022417405.1 | T | 1 (*6*) |
| *Ictidomys tridecemlineatus* | XP_005331769.1 | I | 10 (*7*) |
| *Neomonachus schauinslandi* | XP_021558093.1 | T | 1 (*8*) |
| *Phascolarctos cinereus* | XP_020855496.1 | T | 1(*5*) |
| *Odocoileus virginianus texanus* | XP_020769734.1 | T | 2 (*9*) |
| *Bubalus bubalis* | ARJ58634.1 | T | 1.38 (*1*) |
| *Capra hircus* | NP_001272541.1 | T | 1.5 (*1*) |
| *Equus caballus* | NP_001075305.1 | T | 1 (*1*) |
| *Felis catus* | AFP97799.1 | T | 4 (*10*) |
| *Sarcophilus harrisii* | XP_003773267.1 | T | 17.5 (*5*) |
| *Loxodonta Africana* | XP_003415950.1 | T | 1 (*1*) |
| *Dasypus novemcinctus* | XP_004466930.1 | M | 4 (*11*) |
| *Pteropus vampyrus* | XP_011358956.1 | T | 1 (*12*) |
| *Myotis lucifugus* | XP_006095704.1 | T | 1 (*13*) |
| *Physeter catodon* | XP_007108070.1 | T | 1 (*14*) |
| *Desmodus rotundus* | XP_024433366.1 | T | 1 (*15*) |
| *Neophocaena asiaeorientalis* | XP_024603511.1 | T | 1 (*16*) |
| *Pteropus alecto* | XP_006916576.1 | T | 1.2 (*17*) |
| *Canis lupus dingo* | XP_025291626.1 | T | 4.5 (*18*) |
| *Oryctolagus cuniculus* | AAV48838.1 | I | 5 (*1*) |
| *Rattus norvegicus* | NP_112300.2 | V | 9 (*19*) |
| *Mus musculus* | Q01776.1 | V | 7.4 |
| *Heterocephalus glaber* | XP_004849709.1 | I | 11 (*20*) |
| *Mus caroli* | XP_021019370.1 | V | 6 (*21*) |
| *Mus pahari* | XP_021066838.1 | V | 6 (*22*) |
| *Papio anubis* | XP_003898885.1 | T | 1 (*23*) |
| *Carlito syrichta* | XP_008069339.1 | T | 1 (*24*) |
| *Aotus nancymaae* | XP_012328857.1 | T | 1 (*25*) |
| *Microcebus murinus* | XP_012616420.1 | T | 2 (*26*) |
| *Theropithecus gelada* | XP_025242022.1 | T | 1 (*1*) |
| *Pan paniscus* | XP_003815858.1 | T | 1 (*1*) |
| *Macaca nemestrina* | XP_011709114.1 | T | 1 (*1*) |
| *Pan troglodytes* | XP_526608.1 | T | 1 (*1*) |
| *Pongo abelii* | XP_024101999.1 | T | 1 (*27*) |
| *Piliocolobus tephrosceles* | XP_023087932.1 | T | *1 (*28*) |
| *Otolemur garnettii* | XP_003801017.1 | A | 1( *29*) |
| *Homo sapiens* | AAB26287.1 | T | 1 (*23*) |

### Supplementary References.


1. J. Werner, E. M. Griebeler, Reproductive Biology and Its Impact on Body Size: Comparative Analysis of Mammalian, Avian and Dinosaurian Reproduction. *PLoS ONE*. **6**, e28442 (2011).
2. K. S. Borge, R. Tønnessen, A. Nødtvedt, A. Indrebø, Litter size at birth in purebred dogs-A retrospective study of 224 breeds. *Theriogenology*. **75**, 911–919 (2011).
3. C. Li, G. Wiener, H. JianLin, L. RuiJun, in *The Yak* (Food and Agriculture Organization of the United Nations Regional Office for Asia and the Pacific, Bangkok, ed. 2, 2003; http://www.fao.org/3/AD347E/ad347e00.htm#Contents).
4. G. W. Wood, R. H. Barrett, Status of Wild Pigs in the United States. *Wildlife Society Bulletin*. **7**, 237–246 (1979).
5. R. M. Nowak, *Walker's Mammals of the World: Monotremes, Marsupials, Afrotherians, Xenarthrans, and Sundatherians* (Johns Hopkins University Press, 2018), *Walker's Mammals*.
6. Beluga (Delphinapterus leucas) longevity, ageing, and life history. *AnAge: the Animal Ageing and Longevity Database*, (available at http://genomics.senescence.info/species/entry.php?species=Delphinapterus_leucas).
7. E. C. Cleary, S. R. Craven, in *Prevention and Control of Wildlife Damage*, S. E. Hygnstrom, R. M. Timm, G. E. Larson, Eds. (University of Nebraska-Lincoln, 1994; https://digitalcommons.unl.edu/icwdmhandbook/).
8. W. F. Perrin, J. G. M. Thewissen, B. Würsig, *Encyclopedia of Marine Mammals* (Elsevier Science & Technology, ed. 2, 2008).
9. R. J. Innes, Odocoileus virginianus. *Fire Effects Information System* (2013), (available at https://www.fs.fed.us/database/feis/animals/mammal/odvi/all.html#ReproductionAndDevelopment).
10. M. v. Root Kustritz, Clinical management of pregnancy in cats. *Theriogenology*. **66**, 145–150 (2006).
11. D. W. Hawthorne, in *Prevention and Control of Wildlife Damage*, S. E. Hygnstrom, R. M. Timm, G. E. Larson, Eds. (University of Nebraska-Lincoln, 1994; https://digitalcommons.unl.edu/icwdmhandbook/).
12. T. H. Kunz, W. R. Hood, L. Nadolny, in *Bats in Captivity*, S. M. Barnard, Ed. (Logos Press, 2010; https://www.researchgate.net/publication/216052772), vol. 2, pp. 223–238.
13. Little brown bat (Myotis lucifugus) longevity, ageing, and life history. *AnAge: the Animal Ageing and Longevity Database*, (available at http://genomics.senescence.info/species/entry.php?species=Myotis_lucifugus).
14. P. Langer, The phases of maternal investment in eutherian mammals. *Zoology*. **111**, 148–162 (2008).
15. Vampire bat (Desmodus rotundus) longevity, ageing, and life history. *AnAge: the Animal Ageing and Longevity Database*, (available at http://genomics.senescence.info/species/entry.php?species=Desmodus_rotundus).
16. C. Chapman. Personal communication. (April 15, 2020).
17. Black flying fox (Pteropus alecto) longevity, ageing, and life history. *AnAge: the Animal Ageing and Longevity Database*, (available at http://genomics.senescence.info/species/entry.php?species=Pteropus_alecto).
18. R. Hudson, H. G. Rödel, M. T. Elizalde, L. Arteaga, G. A. Kennedy, B. P. Smith, Pattern of nipple use by puppies: A comparison of the dingo (Canis dingo) and the domestic dog (Canis familiaris). *Journal of Comparative Psychology*. **130**, 269–277 (2016).
19. R. M. Timm, in *Prevention and Control of Wildlife Damage*, S. E. Hygnstrom, R. M. Timm, G. E. Larson, Eds. (University of Nebraska-Lincoln, 1994; https://digitalcommons.unl.edu/icwdmhandbook/).
20. W. H. Gates, Litter size, birth weight, and early growth rate of mice (Mus musculus). *The Anatomical Record*. **29**, 183–193 (1925).
21. G. D. Zitnik, S. A. Bingel, S. M. Sumi, G. M. Martin, Survival curves, reproductive life span and age-related pathology of Mus caroli. *Laboratory Animal Science*. **42**, 119–126 (1992).
22. V. C. Agrawal, "Taxonomic Studies on Indian Muridae and Hystricidae. Records of the Zoological Survey of India" (Calcutta, 2000), (available at http://faunaofindia.nic.in/PDFVolumes/occpapers/180/index.pdf).
23. P. H. Harvey, T. H. Clutton-Brock, Life History Variation in Primates. *Evolution*. **39**, 559–581 (1985).
24. Philippine tarsier (Carlito syrichta) longevity, ageing, and life history. *AnAge: the Animal Ageing and Longevity Database*, (available at http://genomics.senescence.info/species/entry.php?species=Carlito_syrichta).
25. A. Gozalo, E. Montoya, Reproduction of the owl monkey (Aotus nancymai) (primates:Cebidae) in captivity. *American Journal of Primatology*. **21**, 61–68 (1990).
26. Gray mouse lemur (Microcebus murinus) longevity, ageing, and life history. *AnAge: the Animal Ageing and Longevity Database*, (available at http://genomics.senescence.info/species/entry.php?species=Microcebus_murinus).
27. D. Kelle, D. Fechter, A. Singer, P. Pratje, I. Storch, Determining Sensitive Parameters for the Population Viability of Reintroduced Sumatran Orangutans (Pongo abelii). *International Journal of Primatology*. **34**, 423–442 (2013).
28. D. Wang. Personal communication. (April 15, 2020).
29. Small-eared galago (Otolemur garnettii) longevity, ageing, and life history. *AnAge: the Animal Ageing and Longevity Database*, (available at http://genomics.senescence.info/species/entry.php?species=Otolemur_garnettii).
